# Construction of a macrophage-tropic subtype C HIV-1 mGreenLantern reporter virus for studies on HIV-1 replication and the impact of methamphetamine

**DOI:** 10.1101/2024.10.24.619504

**Authors:** Dina Mofed, Angelo Mandarino, Xuhong Wu, Yuekun Lang, Anjali Gowripalan, Ganjam V. Kalpana, Vinayaka R. Prasad

## Abstract

HIV-1 subtype C viruses are responsible for 50% of global HIV burden. However, nearly all currently available reporter viruses widely used in HIV research are based on subtype B. We constructed and characterized a replication competent HIV-1 subtype C reporter virus expressing mGreenLantern. mGreenLantern sequences were inserted in-frame with *nef* ATG in HIV-1_IndieC1_. As controls, we employed HIV-1_IndieC1_, HIV-1_ADA,_ and HIV-1_NLAD8-GFP-Nef_ viruses. HIV-1_IndieC1-mGreenLantern_ (HIV-1_IndieC1-mGL_) exhibited characteristics of the parental HIV-1_IndieC1_ virus, including its infectivity in TZMbl reporter cells and replication competence in macrophages. To further characterize HIV-1_IndieC1-mGL_ virus, we tested its responsiveness to CCL2 levels, a characteristic feature of subtype B HIV-1 that is missing in subtype C. CCL2 immunodepletion inhibited the production of HIV-1_ADA_ and HIV-1_NLAD8-GFP-Nef_ as expected, but not that of HIV-1_IndieC1-mGL_ as previously reported. We also tested the effect of Methamphetamine, as its effect is mediated by NF-κB and since subtype C viruses carry an additional copy of NFκB. We found that methamphetamine increased the replication of all viruses tested in macrophages, however, its effect was much more robust for HIV-1_IndieC1_ and HIV-1_IndieC1-mGL_. Our studies established that HIV-1_IndieC1-mGL_ retains all the characteristics of the parental HIV-1_IndieC1_ and can be a useful tool for HIV-1 subtype C investigations.

## 1. Introduction

HIV-1 subtype C is responsible for a major share — up to 50% — of the global HIV burden [1,2] and displays unique features among group M subtypes. Subtype B HIV-1, on the other hand is responsible for only about 10% of the global HIV burden. Subtype C HIV-1 is the least fit of all group M HIV-1 subtypes and their circulating recombinant forms (CRFs) [3.4]. HIV-1 subtype C is also less likely to undergo a coreceptor usage switch and hence a greater proportion of viruses in people living with HIV are likely to be monocyte-tropic viruses [5,6]. While most HIV-1 subtypes have two copies of NFκB enhancer, subtype C viruses have three NF-κB binding sites in the long terminal repeat (LTR) leading to higher transcriptional activity *in vitro* and higher viral loads in people living with HIV-1. In fact, reports have shown the appearance of a fourth, novel NFκB variant in HIV-1C viruses in India [7,8]. HIV-1 Gag p6 contains two late motifs — PTAP and LYPX — which recruit the ESCRT factors TSG101 or ALIX respectively to trigger membrane fission at the bud neck and facilitate virus release [9]. Duplication of the PTAP motif is seen rarely in subtype B viruses in infected individuals [10]. In contrast, in HIV-1 subtype C-infected individuals, PTAP duplication is seen in up to 30% of infected individuals [11]. Additionally, a functional LYPX motif is absent in nearly all HIV-1 subtype C HIV-1 due to a LY dipeptide deletion [12,13]. However, it has been reported that in some antiretroviral treatment failure cases, variant viruses emerge containing a PYxE tetrapeptide sequence [14]. Such acquired PYxE insertions restore binding to ALIX, increase virus fitness, and restore increased virus budding in response to CCL2 signaling [12]. Furthermore, while all other HIV-1 subtypes retain a highly conserved dicysteine motif in the Tat protein that is part of a conserved feature between HIV-1 Tat proteins and β-chemokines, HIV −1 subtype C lacks this feature due to a C31S substitution in the dicysteine motif [15].

Genetically engineered reporter HIV-1 constructs have allowed researchers to monitor the kinetics of infection and replication *in vitro*. Over the years, a large number of recombinant reporter viruses have been designed, developed, and used as tools to study various aspects of HIV-1 replication and its neutralization by antibodies. For example, reporter viruses carrying *luciferase* gene are used to monitor virus-cell membrane fusion [16], nuclear entry, loss of capsid integrity, and uncoating [17]. Reporter viruses carrying *luciferase or green fluorescent protein* (GFP) have been used to determine the infectivity of HIV-1 variants, to test their sensitivity to antiretrovirals, studying viral tropism or understanding the immunological responses to infection. Reporter viruses with Nef substitutions are generally used to study early events, while Gag-fusion reporter vectors are designed to study both early and late events of virus assembly and release [18]. Reporter viruses that contained fluorescent proteins fused to Vpr or envelope proteins have also been generated to visualize and measure virion production and release. The Vpr-GFP fusion protein helps monitor HIV-1 virions released *via* fluorescence microscopy or flow cytometry, allowing real-time tracking of viral release and high-throughput screening for factors influencing viral release [19]. The use of Gaussia *luciferase* fusion with HIV-1 envelope protein allows detection of low levels of virion release. Similarly, NanoLuc-containing bioluminescent HIV-1 reporter viruses facilitated high sensitivity measurement of virus particle production. Kirui and Freed [20] showed how the use of this new reporter virus enhances the efficacy of quantification. More recently, dual reporter viruses have been developed to separately monitor the expression of LTR-dependent and LTR-independent reporter genes to monitor the time to latency from the time of infection [21].

The HIV-1 reporter viruses mentioned above are only a fraction of a larger number that have been reported over the years and that are in use currently. It is unusual that currently there is only one reporter virus based on HIV-1 subtype C [33]. Reporter viruses exemplified by the ones mentioned above have helped vastly advance our knowledge of the biology and pathogenesis of HIV-1 subtype B. A similar progress with subtype C viruses is much needed.

In this study, we generated a mGreenLantern-tagged full-length and replication competent HIV-1_IndieC1_ reporter virus clone for studies in myeloid cells. First, we established that the infectivity and replication competence in macrophages of HIV-1_IndieC1-mGreenLantern_ (HIV-1_IndieC1-mGL_) virus was similar to that of the parental virus. Second, we investigated whether the new reporter virus retains a characteristic feature of HIV-1 subtype C viruses, namely, the inability to respond to CCL2-mediated modulation of virus production owing to the characteristic absence of a functional LYPX motif in the Gag p6 late domain. HIV-1 budding is mediated by the cellular endosomal sorting complexes required for transport (ESCRT) machinery, specifically the ESCRT III complex which is made up of charged multivesicular body proteins (CHMPs). Such proteins are required both for the abscission step to allow cells to divide or for viruses such as HIV-1 to bud from the infected cell. CHMPs transiently form long filaments that coil at the site of membrane constriction and trigger membrane cleavage. CHMPs are recruited to the sites of HIV-1 budding at the plasma membrane *via* late domain motifs such as PTAP or LYPX on the Gag polyprotein. It has been previously shown that HIV-1 subtype C isolates lack LYPX and are thus unable to recruit ALIX, an ESCRT III protein adapter protein that enhances budding by recruiting CHMP4b protein. We have previously shown that CCL2 treatment stimulates the mobilization of ALIX from the F-actin cytoskeleton to the cytoplasm making it possible for HIV-1 Gag to recruit ALIX and enhance budding. Due to the lack of LYPX motif on subtype C HIV-1 Gag protein, these viruses are unable to respond to CCL2 in the media by recruiting more ALIX and generating more viruses *via* increased budding. It has also been shown that blocking CCL2-signaling *via* the addition of anti-CCL2, which has been shown to result in a near complete colocalization of ALIX with F-actin structures. We tested the new reporter virus for its ability to respond to CCL2 or anti-CCL2 treatment and found that the production of virus is not modulated by the levels of CCL2 present in the medium. Finally, we tested the responsiveness of the new HIV-1 subtype C reporter to methamphetamine (Meth). We report that HIV-1 subtype C virus replication is enhanced by Meth, and the increase is to a much higher level than that observed in HIV-1 subtype B.

## 1. Materials and Methods

### 2.1. Cells, cell lines and infectious clones of HIV-1

293T cells were obtained from ATCC and TZM-bl cells from the NIH AIDS Reagent Repository. Both cell types were maintained in Dulbecco’s Modified Eagle Medium (DMEM) (Thermofisher Scientific, USA) supplemented with 10% FBS (Thermofisher Scientific, USA) and 1% Penicillin-Streptomycin (Thermofisher Scientific, USA), and incubated at 37 °C with 5% CO_2_. Human monocyte-derived macrophages (MDMs) were obtained by differentiating CD14_+_ monocytes isolated from peripheral blood mononuclear cells (PBMCs) by negative selection. PBMCs were purified from leukopaks from New York Blood Center.

### 2.2 Plasmids

pIndieC1 was a gift from Dr. U. Ranga, pNLAD8-GFPNef was a gift from Dr. J. Karn, pADA was a gift from Keith Peden. The pcDNA 3.1-mGreenLantern and pEF1a-IRES-Neo were from Addgene.

### 2.3. Construction of HIV-1_IndieC1-mGL_ reporter virus

The mGreenLantern-IRES cassette was inserted into the backbone of pIndieC1 by overlap PCR amplification of five DNA fragments. Fragment 1 (2614 bp) represents a part of gp160 starting from Pac I restriction site, fragment 2 represents mGreenLantern (720 bp) from pcDNA3.1-mGreenLantern [22], fragment 3 represents IRES (553 bp) derived from pEF1a-IRES-Neo [23], fragment 4 represents Nef (621 bp), and fragment 5 represents the rest of the pIndieC backbone (3575 bp) which ends at the AatII restriction site. Using the built-in repeat sequences at the joining ends, fragments I and II were combined by overlap PCR, followed by sequentially adding fragments III, IV, and V all *via* overlap PCR as above. All PCR reactions were performed using Q5 High-fidelity DNA polymerase (New England Biolabs). Primers used for PCR reactions are listed in Table 1. The final 8 kb product (Pac1-gp160-mGreenLantern-IRES-Nef-LTR-*AatII*) was digested with Pac I and Aat II and ligated to 6.7kb Pac I-*AatII* fragment derived from parental pIndieC1 plasmid to recreate the entire virus clone with the mGreenLantern reporter gene (Figure 1). The presence of mGreenLantern-IRES in the final pIndieC1-mGreenLantern plasmid and the absence of undesired mutations were validated *via* nanopore (Plasmidsaurus, USA) and Sanger sequencing.

**Table 1:**
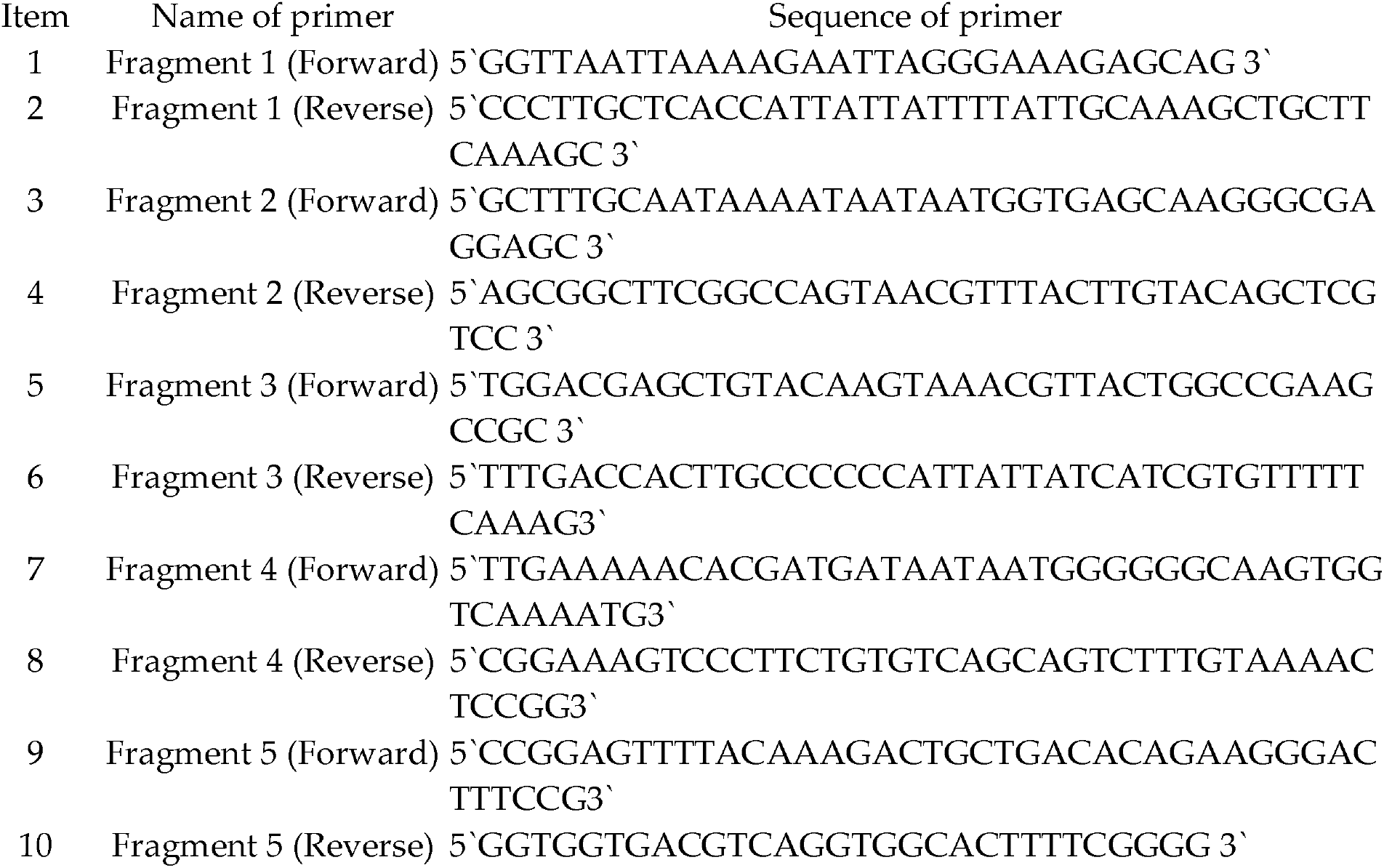
List of primers used in overlap PCR to build the Pac1-gp160-mGreenLantern-IRES-Nef-LTR-AatII fragment.

**Figure 1:**
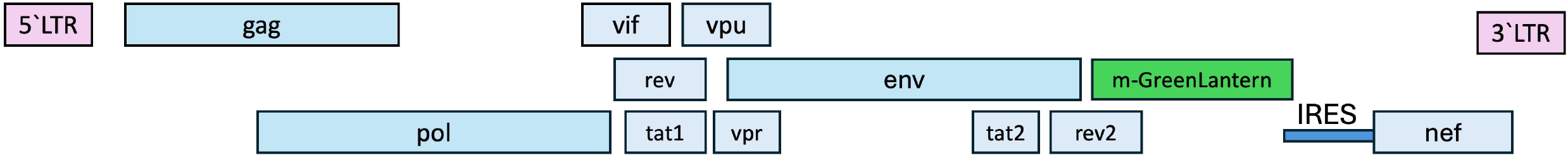
Systematic representation of the constructed mGreenLantern expressing the HIV-1 subtype C reporter virus and the IRES-mGreenlantern cassette inserted in-frame between Env and Nef genes.

### 2.4. Generating high titer virus stocks, and testing their infectivity

CD14_+_ monocytes were plated with Monocytes attachment medium (PromoCell, USA) and differentiated into macrophages by adding DMEM supplemented with 10% FBS, 1% Penicillin-Streptomycin, and rhM-CSF (5 ng/ml, Peprotech), and grown in a cell culture incubator for ten days.

To generate high-titer viruses, plasmid DNAs of infectious molecular clones HIV-1_ADA_ (subtype B) [24], HIV-1_IndieC1_ (subtype C) [25], HIV-1_NLAD8-GFP-Nef_ [26], and HIV-1_IndieC1-mGL_ reporter were transfected separately (20µg each) into 293T cells seeded at a density of 10 x 10^6^ cells in a 10 cm plate using Lipofectamine 3000 (Invitrogen, ThermoFisher, USA). Media containing infectious viruses were collected at 24h post-transfection for p24 measurement. The p24 levels were measured using AlphaLisa detection kit (Perkin Elmer, cat #AL291F) for HIV-1_ADA_ and HIV-1_NLAD8-GFP-Nef_ (subtype B) or using HIV-1 p24 Antigen Capture Assay kit (ABL) for HIV-1_IndieC1_ and HIV-1_IndieC1-mGL_ (subtype C). The infectivity of viruses thus generated was tested using TZM-bl luciferase assay. Briefly, TZM-bl cells were seeded in 12-well plates at a density of 2 x10^5^ cells per well and incubated at 37°C with 5% CO_2_ for 24h. Cells were infected with each of the four HIV-1 viruses mentioned above at increasing multiplicities of infection (MOI) equivalent to 31.25, 62.5, 125, 250, and 500ng p24 per 2 x10^5^ cells and incubated at 37°C with 5% CO_2_ for 24h. Cells were washed twice with 1X PBS and lysed using lysis buffer in the Luciferase assay kit (Promega). For each virus, 50 µl of cell lysate from each MOI was used to measure the luciferase activity.

### 2.5. Multiday HIV-1 replication assay in monocyte-derived macrophages

Macrophage infection and replication of HIV-1_ADA,_ HIV-_1NLAD8-GFP-Nef_ reporter, HIV-1_IndieC1,_ and HIV-1_IndieC1-mGL_ reporter viruses were tested in monocyte-derived macrophages (MDMs). Briefly, CD14_+_ monocytes were seeded in 6-well plates at a density of 1×10^6^ cells per well and differentiated into macrophages as described above. MDMs were infected with each of the 4 different viruses separately at 10ng p24 per 106 cells. Half of the media (1ml) was removed and replenished with fresh medium every three days throughout the multiday replication experiment. Replication competence was determined by measuring p24 in cultured infected media.

### 2.6. Measuring the effect of CCL2 and anti-CCL2 on the replication of HIV-1 isolates

To determine the effect of CCL2 and anti-CCL2 on the replication of HIV-1 (subtype B and subtype C), 1×10^6^ MDMs were infected with HIV-_1NLAD8-GFP-Nef_ and HIV-1_IndieC1-mGL_ reporter viruses at 10ng p24 per 106 cells. Infected MDMs were treated with CCL2 (250 ng/mL), anti-CCL2 (2.5 mg/mL), or an isotype control antibody (2.5 mg/mL) [12]. Half media and all the treatments were replenished every 3 days for 30 days. Replication of HIV-_1NLAD8-GFP-Nef_ and HIV-1_IndieC1-mGL_ reporter viruses was monitored by tracking the levels of cellular green fluorescence using an inverted fluorescence microscope (Zeiss Axio Observer) or by measuring p24 levels in the cultured infected media as mentioned above.

### 2.7. Measuring the effect of methamphetamine on the replication of HIV-1 isolates

To investigate the effect of Meth on the HIV-1 (subtype B and subtype C) replication, MDMs were seeded in 6 well plates at a density of 1 ×10^6^ cells/well, and infected at day 10 with HIV-1_ADA,_ HIV-1_NLAD8-GFP-Nef,_ HIV-1_IndieC1_ or HIV-1_IndieC1-mGL_ viruses respectively at 10ng p24/106 cells. Infected MDMs were treated with Meth at three different concentrations (100, 250, 500µM). Every three days, half media was replenished with new media containing fresh Meth for 15 days. The effect of Meth on the replication of all infected viruses was determined by monitoring p24 production in the media. In addition, the effect on the replication of HIV-_1NLAD8-GFP-Nef_ and HIV-1_IndieC1-mGL_ viruses was investigated by tracking the levels of cellular green fluorescence (Zeiss Axio Observer).

### 2.8. Quantification of fluorescence intensity

For quantification of GFP or mGreenLantern fluorescence intensities in Figures 4B, 4C and 5B, we used NIH Image J. The intensities for each time point were calculated from images taken from three replicates and plotted in Figures 4D and 5C respectively.

### 2.9. Statistical analysis

At least three independent replicates were done for each experiment, and the mean was calculated in all cases. Statistical analyses were performed using GraphPad Prism 8.0 (GraphPad Software, San Diego, CA, USA).

## 3. Results

### 3.1 HIV-1 _IndieC1-mGL_ reporter virus is a replication competent virus

We wished to test the infectivity and replication competence of the new reporter virus HIV-1_IndieC1-mGL_. We prepared high titer virus stocks of HIV-1_IndieC1-mGL_ and its parental HIV-1_IndieC1_ virus. For comparison, we also generated virus stocks of a subtype B reporter virus, HIV-1_NLAD8-GFP-Nef,_ and its parental virus HIV-1_ADA_. First, we tested the infectivity of these four viruses employing the TZM-bl luciferase assay. TZM-bl cells carry an integrated copy of the luciferase reporter gene under the control of the HIV-1 LTR [27], and thus respond with the induction of luciferase activity that corresponds to the degree of infection. TZM-bl cells were infected with increasing virus inputs for each of the four viruses. The results indicated that the infectivity of HIV-1_IndieC1-mGL_ reporter virus was comparable to its parental virus HIV-1_IndieC1_ (Figure 2A). In our comparison of infectivities of subtype B virus (HIV-1_ADA_) and the subtype B reporter virus, HIV-1_NLAD8-GFP-Nef_, the reporter virus was consistently higher than that of HIV-1_ADA_ by about 2-fold at all multiplicities of infection (Figure 2B). This is likely due to the chimeric derivation of this virus [26]. HIV-1_NLAD8-GFP-Nef_ virus is based on HIV-1_NL4-3_ provirus but replaces the *env* gene with that from HIV-1_ADA_. As the only common feature between HIV-1_ADA_ and HIV-1_NLAD8-GFP-Nef_ is the *env* gene of HIV-1_ADA_, the robustness of the virus may be attributable to sequences outside *env*.

**Figure 2:**
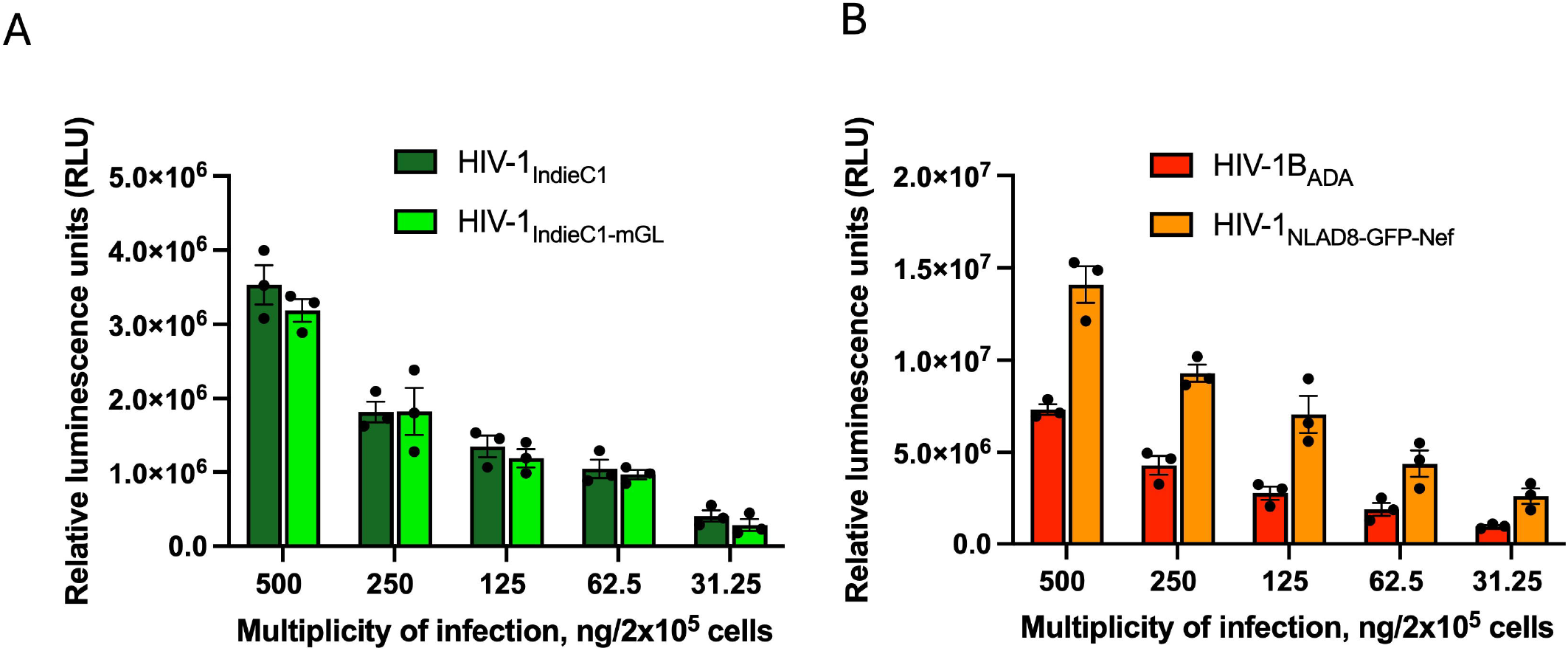
Infectivity of HIV-1 subtype C and B viruses *via* luciferase assay in TZM-bl cells. TZM-bl cells were infected with HIV-1 subtype C and B with different MOI (500, 250, 125, 62.5, and 31.25) ng p24/2 ×10^5^ cells and incubated at 37 °C with 5% CO_2_ for 24hr. A. Infectivity of recombinant HIV-1_IndieC1-mGL_ reporter virus was comparable to the parental HIV-1_IndieC1_. B. Infectivity of recombinant HIV-_1NLAD8-GFP-Nef_ comparable to HIV-1_ADA_. Data are represented as mean ± SEM (n = 3).

Next, we examined the replication competence of the HIV-1 subtype C reporter virus alongside the control viruses (HIV-1_IndieC1_, HIV-1_ADA,_ and HIV-1_NLAD8-GFP-Nef_) in MDMs in a 30-day replication assay. Our results show that HIV-1_IndieC1-mGL_ reporter virus replication in macrophages is comparable to that of its parental HIV-1_IndieC1_ virus (Figure 3A). It is well known that HIV-1 subtype C isolates replicate at a somewhat lower efficiency than subtype B viruses. Accordingly, we find that both HIV-1_ADA_ and HIV-1_NLAD8-GFP-Nef_ viruses replicated at a higher rate reaching a peak virus production that was about 3-fold higher than that of subtype C viruses (compare Figures 3A and 3B). Collectively, the results of the p24 to infectivity ratios and the replication competence experiment show that the HIV-1_IndieC1-mGL_ is a replication competent virus similar to the parental HIV-1_IndieC1_.

**Figure 3:**
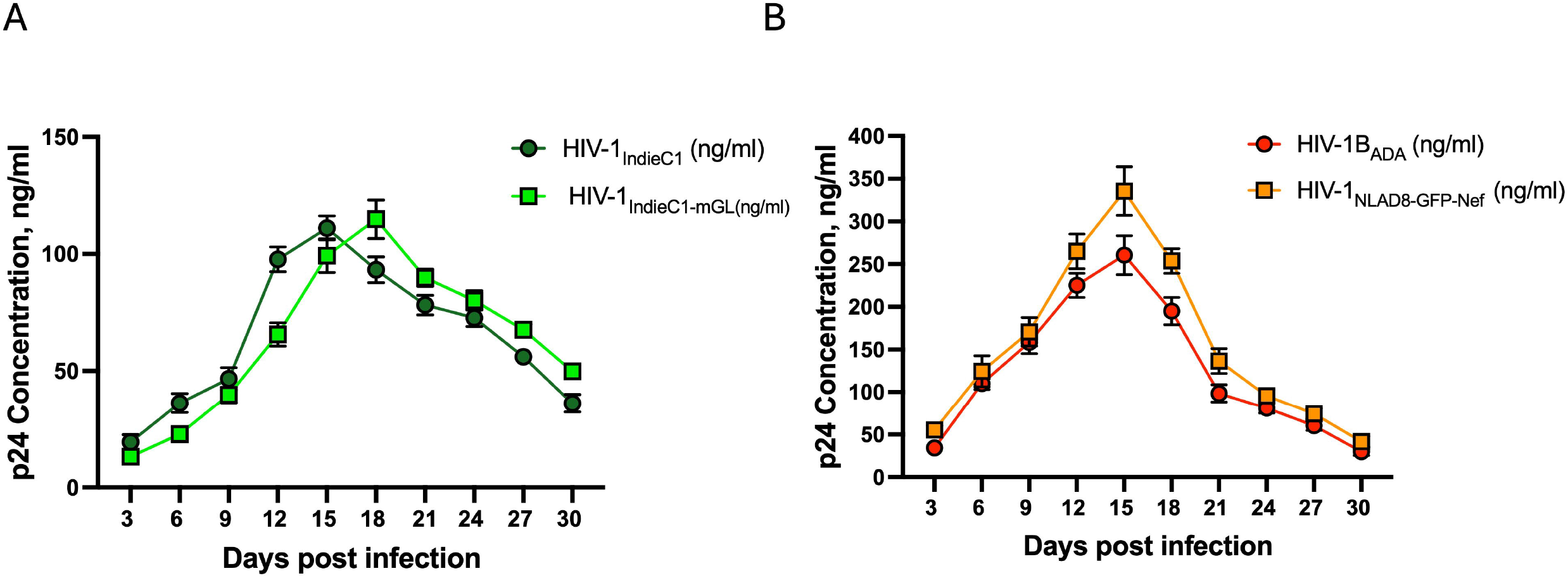
Measurement of replication efficiency of HIV-1 (Subtype B and Subtype C) using a multi-day replication assay in MDMs. MDMs were infected with 10ng/1×10^6^ cells, either subtype C or B. A. Replication competence of recombinant HIV-1_IndieC1-mGL_ reporter virus comparable to the wild-type vector, HIV-1_IndieC1_. B. Replication competence of recombinant HIV-1_NLAD8-GFP-Nef_ comparable to HIV-1_ADA_. The experiment was performed three times with MDMs from three separate donors with two experimental replicates per experiment. Data are represented as mean ± SEM (n = 3).

### 3.2 The production of HIV-1_IndieC1-mGL_ in MDMs is not responsive to CCL2 levels

Previous reports showed that CCL2 immuno-depletion suppresses HIV-1 subtype B replication in macrophages [12, 28] and that the addition of CCL2 enhances virus replication [12]. We showed that this CCL2-dependent modulation of HIV-1 replication was due to the mobilization of ALIX from F-actin stress-fibers to the soluble cytoplasm by CCL2 signaling and this required the presence of late domain motif LYPX in Gag p6, which is responsible for ALIX recruitment to virus assembly and budding sites on the plasma membrane. Furthermore, we reported that 99.8% of the 495 full-length subtype C virus sequences tested lacked a functional LYPX motif due to a highly conserved LY dipeptide deletion in Gag p6. Thus, the subtype C virus HIV-1_IndieC1_ was unresponsive to levels of CCL2 in the media due to its inability to recruit ALIX [12]. Since the efficiency of budding and release of virus particles is not responsive to the levels of CCL2 in HIV-1 subtype C viruses, we tested the HIV-1_IndieC1-mGL_ reporter virus for its CCL2-responsiveness. We infected MDMs with either HIV-1_IndieC1-mGL_ or the parental HIV-1_IndieC1_ virus. For each virus, four separate conditions were employed: untreated, CCL2 treated, anti-CCL2 treated or isotype control IgG treated and the virus particle release measured over a period of 15 days. We also included subtype B controls – HIV-1_ADA_ and HIV-1_NLAD8-GFP-Nef_. As shown in our results, the anti-CCL2 treatment resulted in a substantial reduction (∼6-fold) in the production of subtype B viruses compared to untreated infected MDMs or isotype antibody controls (Figure 4A, Right panel). Similarly, CCL2 treatment enhanced the virus production of subtype B viruses by ∼ 1.8-fold compared to untreated or isotype antibody treated MDMs (Figure 4A, Right panel). In contrast, virus production levels for subtype C viruses, as expected, was unaffected by levels of CCL2 in the medium (Figure 4A, Left Panel).

In addition to monitoring p24 production, we monitored the infection of MDM by HIV-1_IndieC1-mGL_ and HIV-1_NLAD8-GFP-Nef_ viruses by documenting mGreenLantern or GFP fluorescence respectively of the infected cells. For both viruses, we observed a gradual increase in the total fluorescence intensity from the time of infection to day 15 post-infection, indicating peak activity. For HIV-1_NLAD8-GFP-Nef_, treatment with anti-CCL2 led to a considerable reduction in GFP fluorescence compared to untreated or isotype antibody treated, infected MDMs (Figure 4C). Similarly, treatment with CCL2 led to an increase in GFP fluorescence throughout the culture up to day 15 (Figure 4C) as reported previously [12]. In contrast, addition of CCL2, anti-CCL2, or the isotype control antibody did not change the fluorescence intensity of HIV-1_IndieC1-mGL_-infected MDMs (Figure 4B). A fluorescence intensity plot for each treatment for infection with each reporter showed results consistent with the observation that the subtype C reporter is refractory to CCL2 levels unlike the subtype B reporter virus (Figure 4D, compare Left and Right panels). Furthermore, the pattern of responsiveness to CCL2 levels by the two reporter viruses (Figure 4D) was similar to that seen when the p24 production was plotted (Figure 4A). These results together indicated that the replication pattern of the newly created HIV-1_IndieC1-mGL_ follows the patten of the wildtype HIV-1_IndieC1_ and that the replication of this virus can be monitored by measuring p24 as well as by imaging and by following the fluorescence patterns.

**Figure 4:**
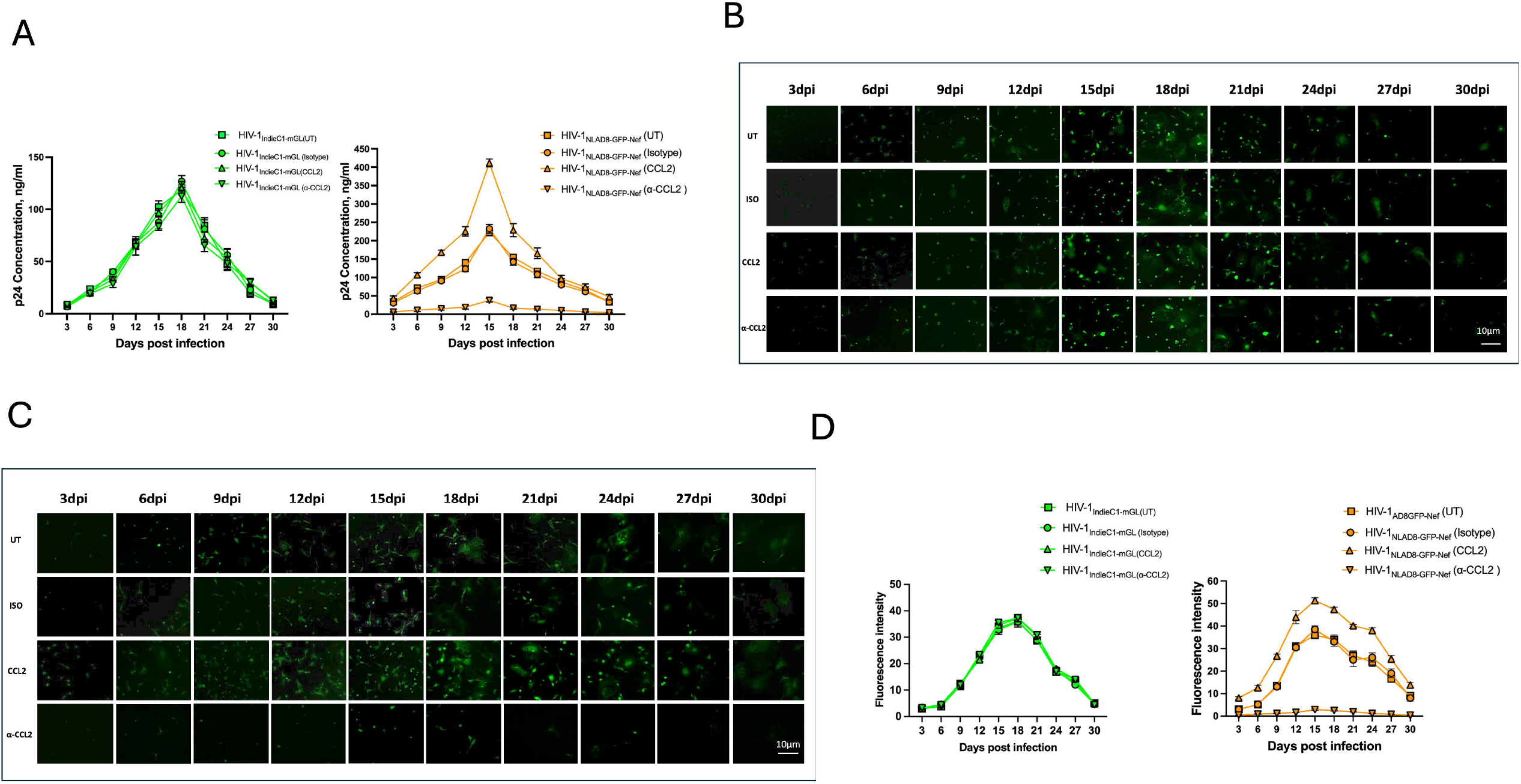
Effect of CCL2 levels on HIV-1 subtype C and B replication in MDMs. A. MDMs were infected with HIV-1_IndieC1-mGL_ reporter virus (Left) or HIV-1_NLAD8-GFP-Nef_ (Right). The infected MDMs were treated with CCL2 (0.25 mg/ml), α-CCL2 (2.5 mg/ ml), or isotype control antibody (2.5 mg/ml). UT refers to no treatment. B. Tracking the effect of CCL2 on HIV-1 _IndieC1-mGL_ reporter replication in MDMs using cellular mGreenLantern. dpi, days post infection. The scale bar for all panels is the same as that shown in the panel corresponding to α-CCL2, 30 dpi. C. Tracking the effect of CCL2 on HIV-1_NLAD8-GFP-Nef_ replication in MDMs using cellular GFP fluorescence intensity. MDMs from three separate donors with two replicates per experiment. Data are represented as mean ± SEM (n = 3). dpi, days post infection. Scale bar for all panels, like in 4C, is shown in the panel corresponding to α-CCL2, 30 dpi. D. Measurement of fluorescence intensity in panels B and C to determine the levels of virus replication for subtype C or subtype B reporter viruses with various treatments.

### 3.3 Methamphetamine pretreatment markedly enhanced HIV-1 subtype C replication in MDMs

Previous studies have reported that Meth increases HIV-1 replication in macrophages [29-31]. Meth is known to trigger the NF-kB signaling pathway in immune cells, including macrophages, and thus its impact on HIV-1 replication is mediated by enhanced LTR transcription [32]. Since HIV-1 subtype C viruses contain 3 NFkB sites, we chose to study the impact of Meth on subtype C replication in MDMs, employing the HIV-1_IndieC1-mGL_ reporter virus. We infected MDMs, in parallel, with HIV-1_ADA_, HIV-1_NLAD8-GFP-Nef,_ HIV-1_IndieC1,_ and HIV-1_IndieC1-mGL_ viruses in the presence or absence of Meth. As indicated by the levels of p24 measured at various times after infection up to day 15, Meth treatment enhanced the replication of all viruses compared to untreated HIV-infected MDMs. However, the increase was more robust for HIV-1 subtype C viruses compared to subtype B viruses (compare Figure 5A top panels to Bottom panels). Also, we noticed that the highest increase in the replication of HIV-1 (subtype B and C) by Meth was observed at a concentration of 100µM, while the increases were progressively lower at 250 and 500 µM respectively. For example, the increase in p24 levels in the presence of Meth over its absence for HIV-1_IndieC-mGL_ was ∼4.5-fold at 100µM, 3.65-fold at 250 µM, and 2.58-fold at 500 µM (Figure 5A, Top right). Increases for HIV-1_IndieC1_ were remarkably similar to that observed for HIV-1_IndieC-mGL_ at each of the three Meth concentrations (Figure 5A, Top left). The increases in virus replication for HIV-1_ADA_ or HIV-1_NLAD8-GFP-Nef_ were significantly lower than subtype C viruses: 3- or 3.8-fold at 100µM, 2.2- or 2.7-fold at 250 µM, and 1.35- or 1.68-fold at 500 µM respectively (Figure 5A Bottom panels).

**Figure 5:**
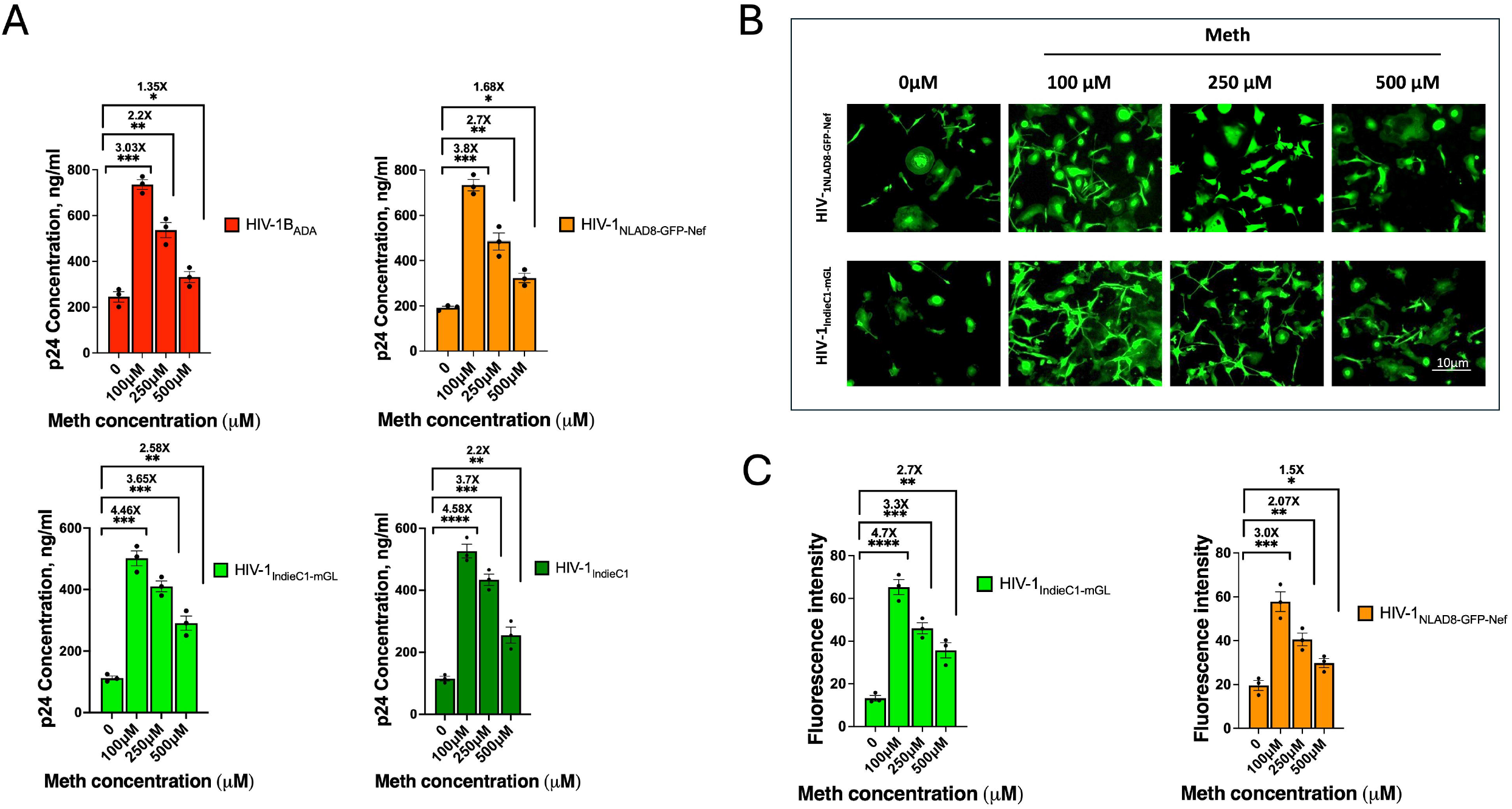
Effect of Meth on HIV-1 replication of various HIV-1 strains tested. A. Differential effect of Meth on HIV-1 subtype C (Top panels) and subtype B (bottom panels) replication in MDMs. MDMs were infected with 10ng/1×10^6^ cells, either subtype C or B viruses. Infected MDMs were treated with Meth at different concentrations (100, 250, and 500µM) or no Meth. The replication of HIV-1_IndieC1_ (Top left); HIV-1_IndieC1-mGL_ reporter (Top right); HIV-1_ADA_ (Bottom left) and HIV-1_NLAD8-GFP-Nef_ (Bottom right) in the presence or absence of Meth as measured *via* p24 concentrations at 15 dpi. P: *, 0.05; **, <0.01; ***, <0.001; and ****, <0.0001. B. Tracking the effect of Meth on HIV-1 _IndieC1-mGL_ or HIV-_1NLAD8-GFP-Nef_ *via* cellular green fluorescence intensity. C. Measurement of fluorescence intensity to determine the effect of Meth on HIV-1 replication from Panel B. MDMs from three separate donors with two replicates per experiment for both HIV-1_IndieC1-mGL_ (Left) and HIV-1_NLAD8-GFP-Nef_ (Right). Data are represented as mean ± SEM (n = 3). Scale bar =10µm. P: *, 0.05; **, <0.01; ***, <0.001; and ****, <0.0001.

To determine if fluorescence intensity of the infected cells reflects the Meth-induced increase in replication of the viruses observed, we also compared the results of cellular green fluorescence intensity in MDMs infected with HIV-1_NLAD8-GFP-Nef_ or HIV-1_IndieC1-mGL_ in the presence or absence of Meth at three different concentrations (100, 250, and 500µM). MDMs infected with either HIV-1_NLAD8-GFP-Nef_ or HIV-1_IndieC1-mGL_ virus displayed an increase in green fluorescence intensity in the presence of Meth compared to its absence (Figure 5B). The increase in fluorescence intensity observed over no Meth was much higher for HIV-1_IndieC1-mGL_ (Figure 5C, Left panel) than for HIV-1_NLAD8-GFP-Nef_ (Figure 5C, Right panel). We note that the fluorescence intensities of GFP and mGreenLantern are intrinsically different. Therefore, the comparisons have been made between treatments with and without Meth but within either HIV-1_IndieC1-mGL_ alone or HIV-1_NLAD8-GFP Nef_ alone. These findings confirm that the impact of Meth on HIV-1 subtype C replication is enhanced to a higher level than subtype B due to the additional NF-kB binding site in HIV-1_IndieC1_ or HIV-1_IndieC1-mGL_.

## 4. Discussion

In this study, we constructed a HIV-1 subtype C virus expressing mGreenLantern *via* the spliced nef mRNA. mGreenLantern is a GFP derivative that has been optimized for efficient folding and rapid expression [22]. As a result, mGreenLantern is 6-times brighter than EGFP. As the reporter virus was based on the macrophage-tropic HIV-1_IndieC1_, this virus would be useful for studying HIV-1 replication in myeloid cells (monocytes, macrophages, and microglia). This new reporter virus allows one to track HIV-1 replication by monitoring mGreenLantern expression in the target cells. This method provides a straightforward visual indicator of infection, allowing for efficient tracking and analysis of viral replication *in vitro*. As shown by our results, HIV-1_IndieC1-mGL_ replicated with equal efficiency to parental HIV-1_IndieC1_ virus — including in the infectivity assay using TZM-bl cells and a multiday replication assay in MDMs (Figure 2 and Figure 3). Not only that HIV-1_IndieC1-mGL_ replicated with equal efficiency to the parental virus, but it also retained its lack of responsiveness to CCL2, which indicates that it also retains other characteristics of HIV-1_indieC1_. HIV-1_IndieC1-mGL_ macrophage-tropic subtype C reporter virus complements the only other currently available subtype C reporter virus by Rai et al, which employed EGFP as a reporter and was based on K3016, a South African subtype C virus that could infect T-cells and PBMCs [33]. Thus, HIV-1_IndieC1-mGL_ represents the only HIV-1_IndieC1_ derived virus with a fluorescence reporter.

Meth is known to contribute to the progression of AIDS by enhancing viral replication rates and the consequent ineffectiveness of antiretrovirals [34]. *In vitro*, Meth increases viral replication in monocytes [31, 35], and in CD4 T cells [29], which can lead to increased viral load in HIV-1-positive, active drug users. However, it must be emphasized that the increase of HIV-1 replication in CD4 T cells [29] is controversial as an earlier study showed that Meth inhibits of HIV-1 replication in T-cells [36]. Interestingly, both these studies have used comparable Meth concentrations in activated CD4 T cells and used the same strain of HIV-1 (HIV-1_BAL_). Bosso et al. showed that the increase in HIV-1 replication observed in the presence of Meth is mediated by NF-kB enhancers in HIV-1 subtype B [37]. The presence of three NF-κB binding sites in the LTR region of subtype C HIV-1 compared to only two NF-κB binding sites in most subtypes of HIV-1, including subtype B, suggested that HIV-1 subtype C may respond to METH with a greater responsiveness than HIV-1 subtype B [37]. Therefore, we examined the effect of Meth treatment on MDMs infected with recombinant HIV-1_IndieC1-mGL_ and HIV-1_NLAD8-GFP-Nef_ viruses. Our results revealed that the Meth treatment of MDMs infected with the HIV-1_IndieC1-mGL_ reporter virus markedly increased the viral replication as compared to that of the infected cultures that were left untreated. This increase was also observed in the parental virus HIV-1_IndieC1_. This increase was higher than that observed for HIV-1_NLAD8-GFP-Nef_ compared to similarly infected untreated MDMs. As expected, Meth treatment of MDMs infected with the parental subtype B virus, HIV-1_ADA_ was comparable to that of HIV-1_NLAD8-GFP-Nef_.

Our results also indicate a dose-dependent response of METH in inducing HIV-1 replication. We find that lower concentrations of METH increase virus replication better. Previous reports have shown increases in HIV-1 replication upon Meth addition. One report studied the effect of Meth on the HIV-infected macrophages at a range of concentrations (1µM to 250 µM) [30]. Quantitation of HIV-1 RNA revealed that the highest increase of HIV by Meth was at 250 µM at 8 days post-infection, an approximately two-fold increase over untreated, infected macrophages. In our experiments for HIV-1 subtype B virus, the highest increases in virus replication were observed at 100µM Meth (up to 3-fold higher) at 15 days post-infection. At the higher concentrations of Meth tested (250µM and 500µm), the extent of inhibition was reduced to 2-fold or less. We do not know the basis for this type of a dose response. Another study investigated the effect of Meth on the expression of the HIV genome in human microglia *in vitro* [32]. They found that Meth treatment increased the transcriptional activity of the HIV-1 long terminal repeat promoter, in a dose-dependent manner. In that study, Meth increased the activity and movement of the transcription factor NF-κB, which is a key player in transcriptional activation from HIV-1 LTR. Blocking NF-κB signaling prevented the Meth-induced activation of HIV-1 LTR. These findings suggested that Meth can directly stimulate HIV gene expression in microglial cells, the primary target cells for HIV-1 in the central nervous system, through NF-κB activation. These results support our finding of a much higher increase in the stimulation of viral replication at 100µM Meth in the presence of 3 NF-κB sites in the LTR as in HIV-1 subtype C HIV-1.

Our mGreenLantern-tagged HIV-1_IndieC1_ reporter virus can be beneficial for *in vitro* studies involving monocytes, microglia, and macrophages. The fact that the reporter virus has all the replication characteristics of its parental clone indicates that it can possibly be also used in small animal models such as humanized mice. Using this reporter virus, we were able to detect the effects of Meth on HIV-1 replication of both subtype B and C in infected MDMs, where the responses to Meth of HIV-1 subtype C were higher than that of HIV-1 subtype B.

## 5. Conclusions

We have shown that a mGreenLantern-tagged subtype C virus, that we constructed, is replication competent, retains characteristic features of subtype C that were tested, and that its responsiveness to Meth treatment is more robust than subtype B. The infectious full-length recombinant mGreenLantern tagged HIV-1_IndieC1_ reporter virus can be an essential tool for understanding the intracellular pathways inside the cells that control the virus replication. Furthermore, it can be used for conducting HIV subtype C-specific research and investigating the viral and host factors influencing the rapidly expanding subtype C infections worldwide.

## Author Contributions

Conceptualization, AM and VP; investigation, DM, YL, AG and XW; writing— original draft preparation, DM and VP; writing—review and editing, AM, YL, AG, GK and VP.; supervision, VP and GK; project administration, VP; funding acquisition, VP. All authors have read and agreed to the published version of the manuscript.

## Data Availability Statement

Following publication, the authors are willing to share all raw data that have been analyzed in the manuscript, and they will be made available to other researchers.

## Acknowledgments

The authors would like to thank Dr. Jonathan Karn for providing the plasmid of HIV-1_NLAD8-GFP-Nef_. In addition, the authors wish to thank Dr. Nosanchuk for METH storage. The work reported here was supported by NIH grant R01.

